# A method to increase the imaging efficiency of tiling light sheet microscopy using scanning non-coaxial beam arrays synchronized with regional virtual confocal slits

**DOI:** 10.1101/2024.08.13.607820

**Authors:** Liang Gao

## Abstract

We present a novel method to improve the imaging efficiency of tiling light sheet microscopy. In the method, scanning non-coaxial beam arrays synchronized with regional virtual confocal slits are used to illuminate imaging plane. There are two advantages. One is the imaging efficiency increases proportional to the number of excitation beams within the non-coaxial beam array. The other is the width of the regional virtual confocal slits could be very wide without admitting off-focus fluorescence generated by the non-coaxial beam array, which makes the method easy to adopt and very robust in practice. We describe the method in detail, characterize the method via numerical simulations. The results suggest that the imaging efficiency and feasibility of the tiling light sheet microscopy could be improved significantly without affecting the 3D imaging ability by using the method. In additions, we propose several configurations to implement the method in practice.

## 1. Introduction

The 3D imaging ability of selective plane illumination microscopy (SPIM), i.e. light sheet microscopy (LSM) relies on the intensity profile of the excitation light sheet. The thickness, light confinement ability and size of the excitation light sheet determine the axial resolution, optical sectioning ability and the size of the field of view (FOV) respectively [1,2]. Thus, confining the illumination light both near the imaging plane as much as possible and along the light propagation direction as far as possible at the same time is required to achieve a high 3D imaging ability over a large FOV in LSM.

Despite the efforts spent to optimize the intensity profile of the excitation light sheet in LSM [3-7], the diffraction of light makes it impossible to reduce the light sheet thickness, improve the light confinement ability and increase the light sheet size at the same time. Therefore, methods other than optimizing the light sheet intensity profile were developed to solve the problem [8-14]. An effective strategy is to move the excitation light sheet quickly along the light propagation direction within the imaging plane, so that both high spatial resolution and good optical sectioning ability can be achieved in a FOV much larger than the size of the light sheet itself [15-23]. Tiling light sheet microscopy (TLSM) uses this strategy to improve the 3D imaging ability of LSM for imaging large samples [19,20].

In TLSM, a large imaging plane is imaged by tiling a short but thin light sheet at multiple positions within the imaging plane and taking an image at each tiling position. The final image is reconstructed using the raw images collected at all tiling positions. It has been demonstrated that TLSM has much better 3D imaging ability than conventional LSM in imaging large samples, ranging from live embryonic specimens to optically cleared biological tissues [20-25]. In addition, TLSM allows adjusting the intensity profile and tiling positions of the excitation light sheet and correcting the excitation light sheet alignment errors easily, so that the 3D imaging ability of TLSM can be optimized in real-time based on the sample and imaging proposes in various applications. Despite the advanced 3D imaging ability of TLSM, the extra camera exposures required by TLSM cause a problem. The imaging speed decreases, and the raw data size increases proportionally to the number of tiles, i.e. the number of camera exposures required per image plane. The problem is troubling when both high 3D imaging ability and high imaging throughput are both required for 3D imaging of large samples using TLSM.

In our previous research, we developed a method using discontinuous light sheets to improve the imaging efficiency of TLSM without sacrificing its 3D imaging ability. In the method, discontinuous light sheets created by scanning coaxial beam arrays synchronized with a virtual confocal slit controlled by the rolling shutter of the detection camera are used to illuminate the imaging plane [26]. We showed that the use of discontinuous light sheets improves the imaging efficiency and reduce the raw data size of TLSM by imaging more effective areas of the imaging plane at each tile without affecting the spatial resolution and optical sectioning ability.

However, the optical alignment is very critical to use discontinuous light sheets created by scanning coaxial beam arrays for sample illumination in TLSM, as the synchronization of the scanning coaxial beam array with the virtual confocal slit on the detection camera that is only several microns wide must always be ensured. In the applications of discontinuous light sheets for imaging large cleared biological tissues, we found it often difficult to satisfy the synchronization requirement for several reasons. First, the synchronization is very sensitive to microscope misalignment. Any misalignment could break the synchronization. Second, the samples are not always perfectly transparent or uniform. The optical aberrations introduced by the sample could also violate the synchronization and reduce the imaging quality. Finally, it is particularly challenging when high spatial resolutions are desired, which often requires the synchronization of the scanning coaxial beam array and a narrow virtual confocal slit of only a few microns wide that is corresponding to several pixel rows on the detection camera. Therefore, we decided to seek a better solution.

Here, we present a method to both improve the imaging efficiency of TLSM and relax the synchronization requirement for using discontinuous light sheets in TLSM. In the method, the exposure of the detection camera is controlled by multiple regional virtual confocal slits (Figure. 1(a)) instead of a global virtual confocal slit as that in regular sCMOS cameras (Figure 1(b)). A discontinuous light sheet, created by scanning a non-coaxial beam array instead of a coaxial beam array is used to illumination the imaging plane. Each beam within the non-coaxial beam array is synchronized with one of the regional virtual confocal slits during imaging. As both excitation beams within the beam array and the regional virtual confocal slits are separated along scanning direction, the width of the regional virtual confocal slits can be much wider than that of the global virtual confocal slit when scanning coaxial beam arrays are used to illuminate the imaging plane without admitting extra off-focus fluorescence background. Therefore, the synchronization could be much easier to achieve and maintain in practice. We describe the method in detail, characterize the method via numerical simulations and propose multiple configurations to implement the method in practice.

**Figure 1.**
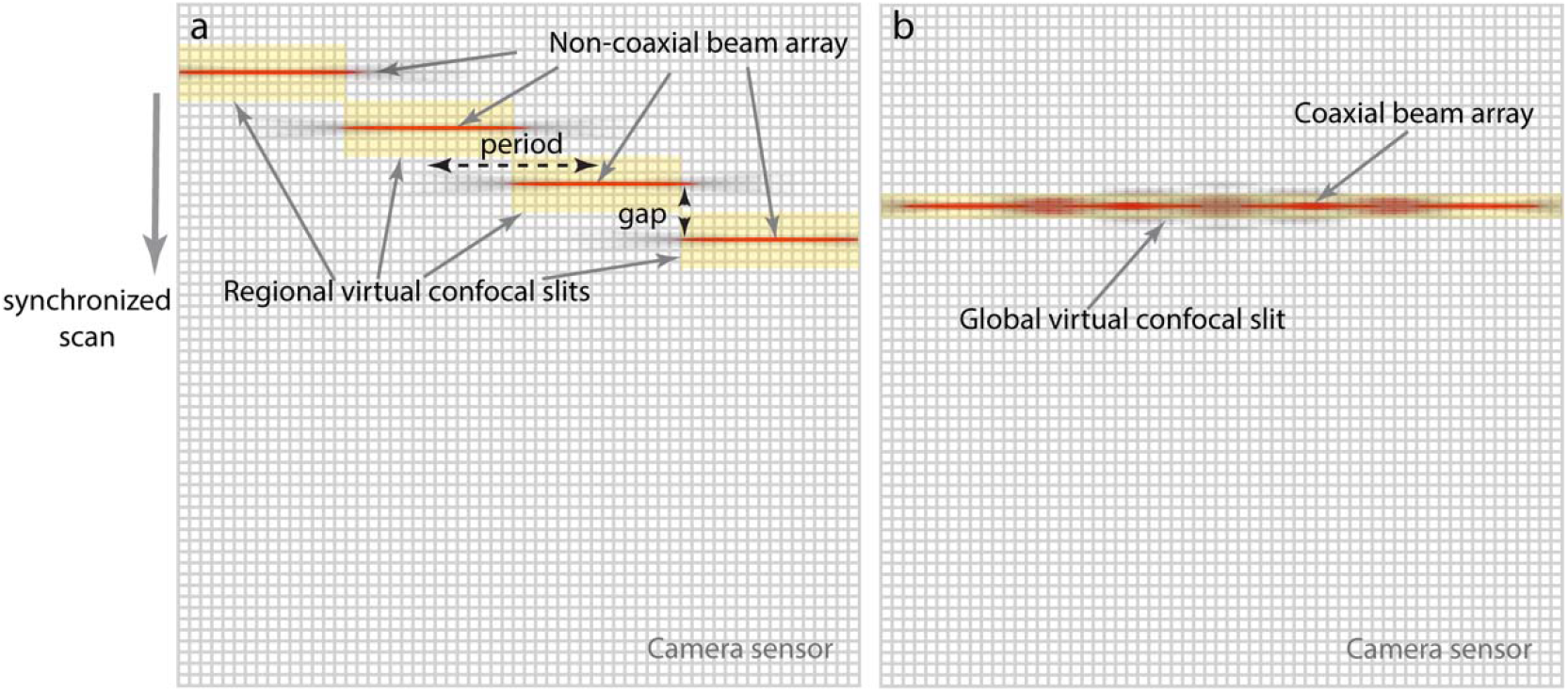
The working mechanism of the method. (a) Illuminating the imaging plane using a scanning non-coaxial beam array synchronized with multiple regional virtual confocal slits. (b) Illuminating the imaging plane using a scanning coaxial beam array synchronized with a global virtual confocal slit. The slit width in (a) can be much wider than that in (b) without admitting extra off-focus fluorescence background, thus allowing much easier and more robust implementation in practice.

## 2. Methods

As described in our previous publication [26], to use discontinuous light sheets created by scanning coaxial beam arrays in TLSM for higher imaging efficiency, the scanning coaxial beam array must be synchronized with a virtual confocal slit that is narrow enough to block the off-focus fluorescence background generated by the coaxial beam array, in which the virtual confocal slit is controlled by the rolling shutter of the sCMOS detection camera (Figure 1(b)). Apparently, the synchronization requirement can be relaxed through two approaches. One is to reduce the crosstalk and off-focus fluorescence of the scanning beam array, and the other is to modify the detection strategy so that the width of the virtual confocal slit can be wider without admitting additional off-focus fluorescence. In light of the consideration, we propose the method of using scanning non-coaxial beam arrays synchronized with regional virtual confocal slits for sample illumination as a solution (Figure 1(a)). The working mechanism is as the following.

First, a non-coaxial beam array instead of a coaxial beam array is scanned to illuminate the imaging plane, in which the excitation beams within the non-coaxial beam array locate in the same imaging plane but separated along the scanning direction. As shown, the separation of the excitation beams along the scanning direction can not only reduce the crosstalk between them but also enable them to stay closer along the excitation light propagation direction due to the reduced crosstalk. In consequence, non-coaxial beam arrays could contain more excitation beams and have less crosstalk between the excitation beams at the same time in comparison to coaxial beam arrays, so that a higher imaging efficiency could be achieved.

Second, as a non-coaxial beam array must be scanned to illuminate the image plane, the off-focus fluorescence generated by the beam array also accumulates and creates a strong off-focus background as all other scanning beams and beam arrays if the detection camera remains exposing during the entire beam array scanning process. Clearly, the detection camera must be synchronized with the non-coaxial beam array scanning to get rid of the off-focus fluorescence background. Instead of using a global virtual confocal slit, we propose using multiple regional virtual confocal slits to match the non-coaxial beam array and reject the off-focus fluorescence created by the excitation beams within the beam array. There are two considerations. First, the excitation beams within a non-coaxial beam array don’t align, so that it is natural to use multiple virtual confocal slits matching the intensity profile of the non-coaxial beam array to block the off-focus fluorescence efficiently without affecting the in-focus fluorescence. Second, as the excitation beams in a non-coaxial beam array are separated along the scanning direction, it is therefore likely to increase the width of the regional virtual confocal slits without admitting additional off-focus fluorescence.

Finally, regional virtual confocal slits are controlled similarly as that of a global virtual confocal slit is controlled by the rolling shutter of sCMOS cameras. For instance, the width, exposure initiation, exposure termination and shifting speed of each regional confocal slit need to be adjusted according to the scanning speed and the gap distance of the non-coaxial beam array, so that the sweeping of the slits, corresponding to the exposure of the pixel rows in different regions of the detection camera, can be synchronized with the scanning of the non-coaxial beam array.

## 3. Results

We characterized the proposed method via numerical simulations. Non-coaxial beam array can be generated using a spatial light modulator (SLM) conjugated to the rear pupil of the excitation objective in TLS microscopes (Figure 2), in which the SLM can be either a continuous phase SLM or a binary phase SLM. The phase maps applied to the SLM can be calculated using a pupil segmentation method described in our previous publication [26].

**Figure 2.**
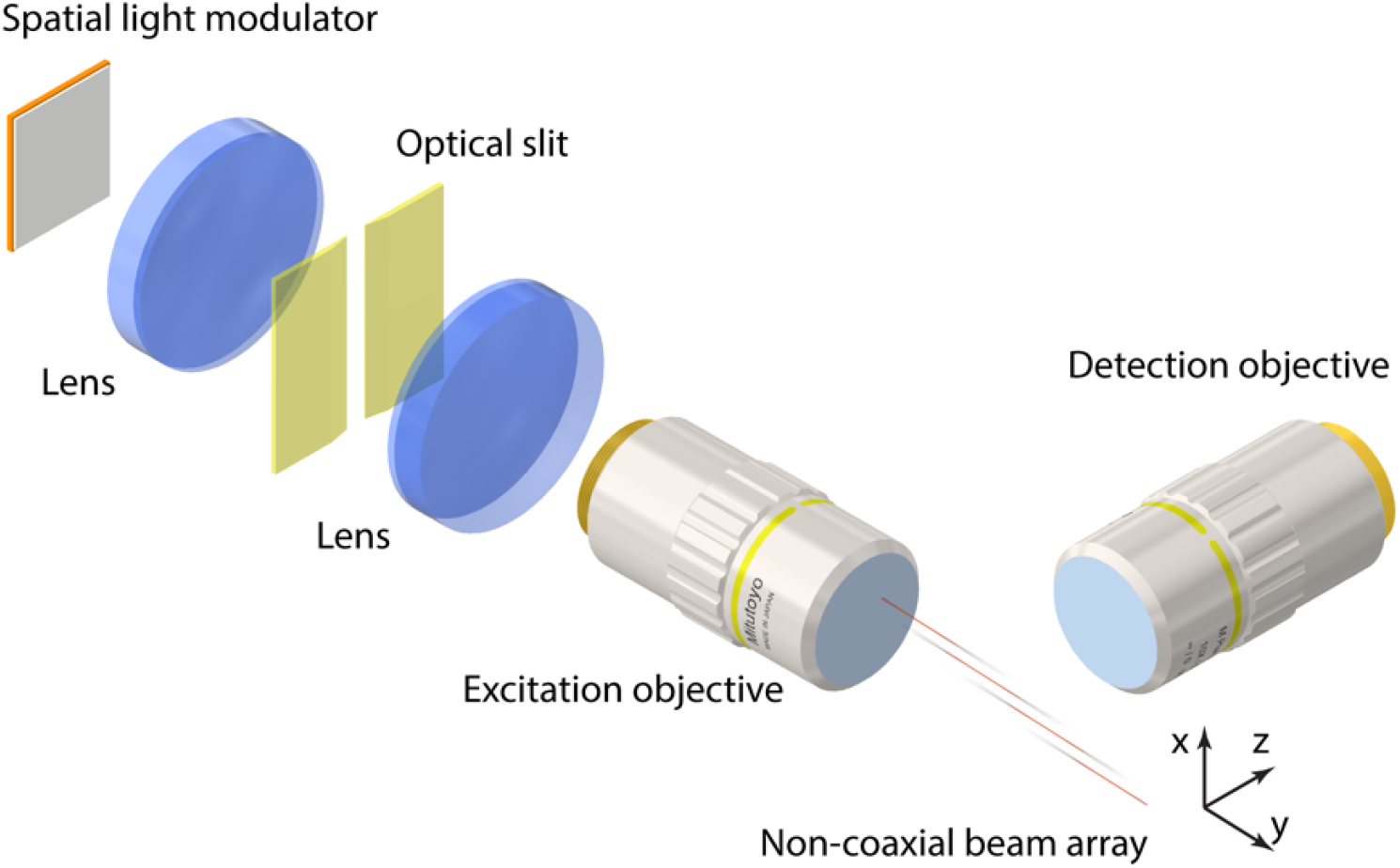
A simplified optical configuration of the TLS microscope showing the generation of noncoaxial beam arrays using a SLM conjugated to the rear pupil of the excitation objective, in which the SLM can be either continuous or binary.

Briefly, the rear pupil of the excitation objective is divided to multiple groups of evenly distributed segments through the SLM conjugated to the rear pupil of the excitation objective. To obtain a non-coaxial beam array, the phased map applied to the SLM is obtained by combining multiple phase maps applied to different groups of pupil segments, so that each group of pupil segments generate one of the excitation beams within a non-coaxial beam array. As shown in Figure 3, a four-beam non-coaxial beam array is generated by dividing the rear pupil of the excitation objective to four groups of evenly distributed radial segments (Figure 3(a)). Four phase maps, which can be either continuous phase maps (Figure 3(b)) or binary phase maps (Figure 3(c)), are combined into one phase map (Figure 3(d) and 3(e)) and applied to the SLM to obtain a four-beam non-coaxial beam array (Figure 3(f) and 3(g)). Furthermore, the properties of the generated non-coaxial beam array, including the excitation beam intensity profile, tiling position, beam array period, beam array gap distance and the number of excitation beams (Figure 4) can be adjusted by applying different phase maps calculated using the described pupil segmentation method. On the other hand, the excitation objective pupil can be divided differently to segments of different shapes, as long as the divided pupil segments distribute evenly.

**Figure 3.**
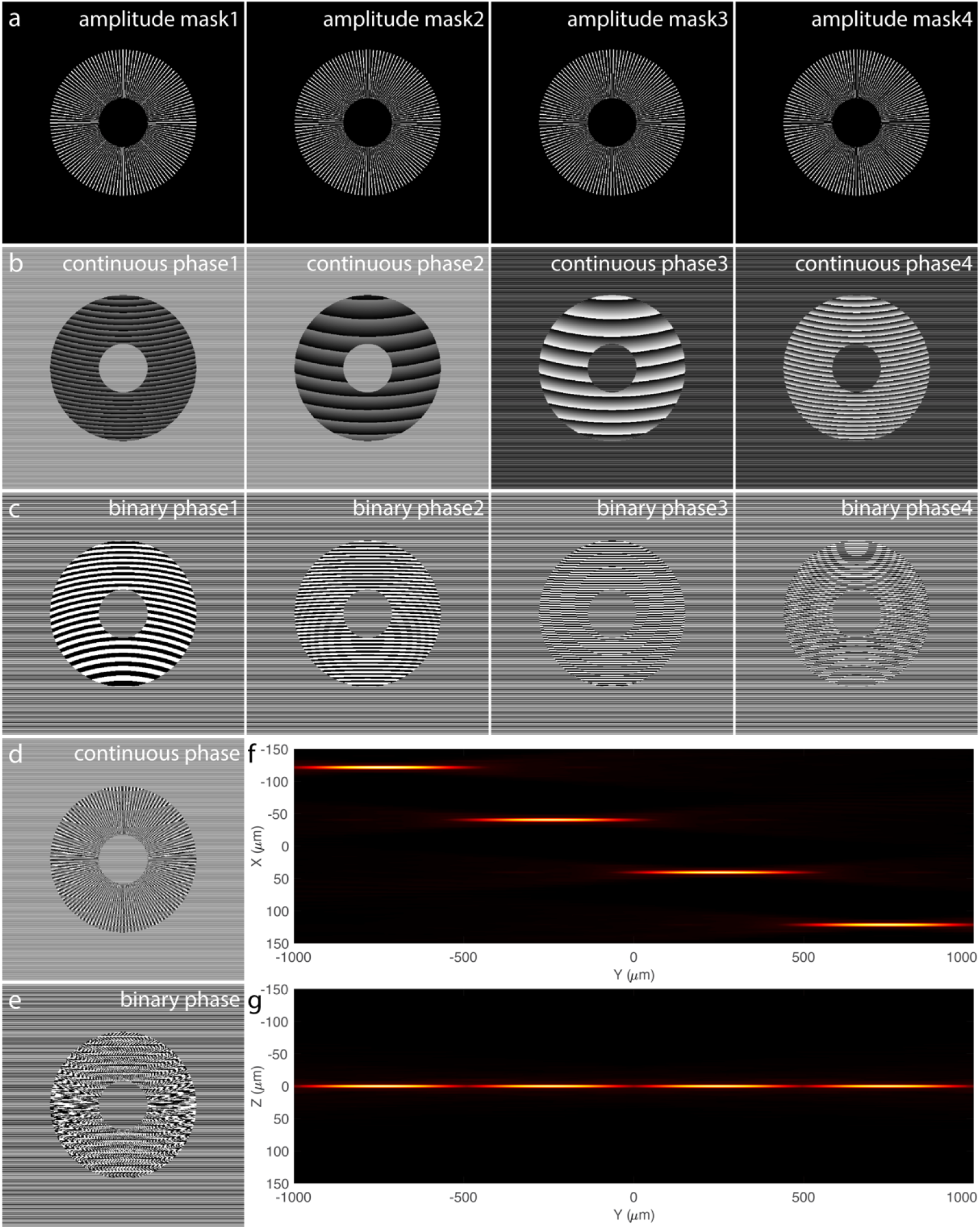
The generation of non-coaxial beam arrays and the calculation of the corresponding phase map using a pupil segmentation method. (a) The excitation pupil is divided to four groups of pupil segments. (b) The continuous phase maps applied to the four groups of pupil segments. (c) The binary phase maps applied to the four groups of pupil segments. (d) The combined continuous phase map used to generate the four-beam non-coaxial beam array in (f) and (g). (e) The combined binary phase map used to generate the four-beam non-coaxial beam array in (f) and (g). (f, g) XY and YZ maximal intensity projections (MIP) of the four-beam non-coaxial beam array generated by applying the continuous phase map in (d) to a continuous phase SLM or the binary phase map in (e) to a binary phase SLM. Excitation numerical aperture (NA): NA_od_=0.06, NA_id_=0.02.

**Figure 4.**
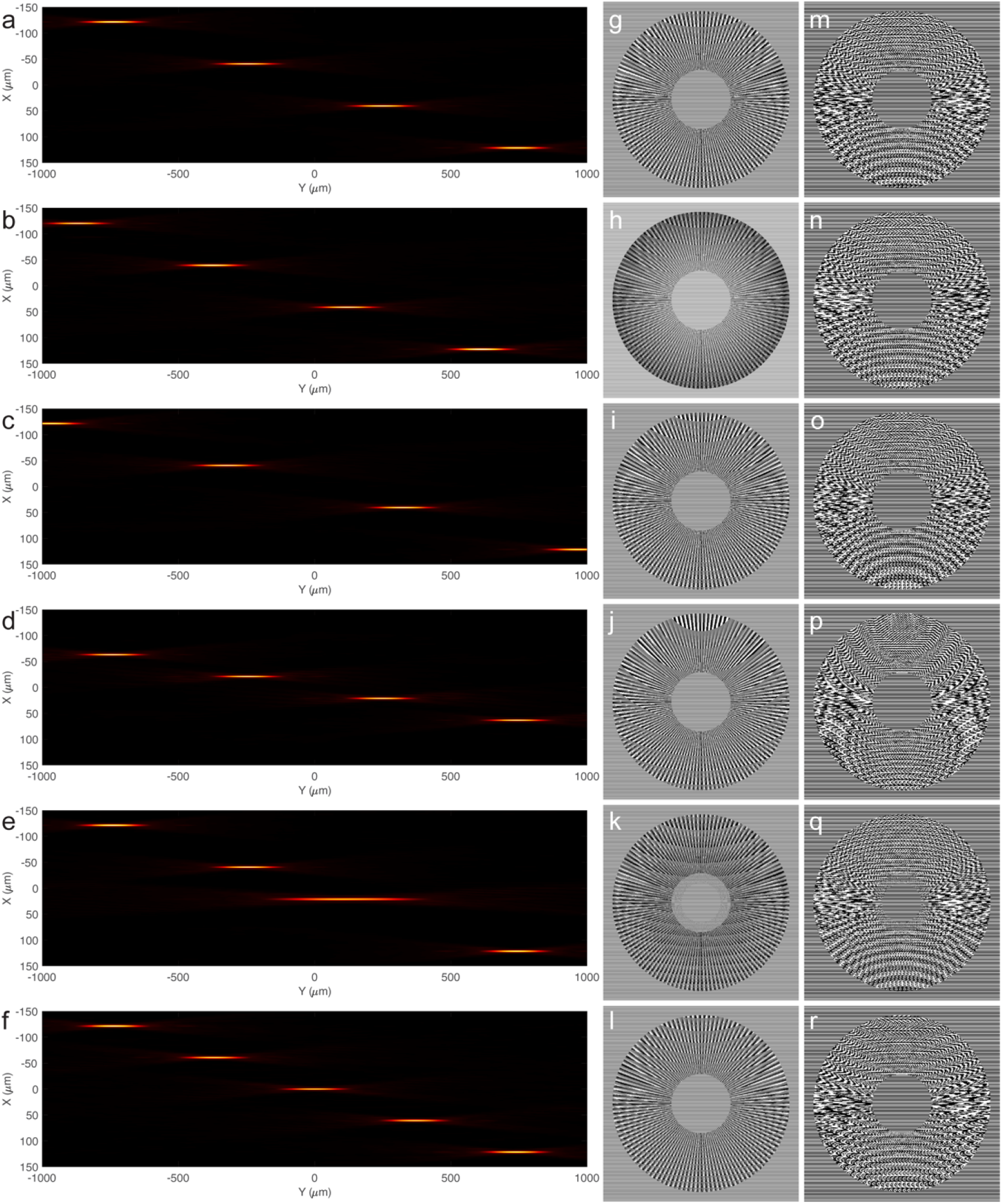
The generation of non-coaxial beam arrays of various properties. Non-coaxial beam arrays of different properties, including the (a) intensity profile of all beams, (b) tiling position, (c) gap distance, (d) beam period, (e) beam number and (f) the properties of individual beams, can be adjusted by applying the continuous phase maps in (g-l) to a continuous phase SLM or binary phase maps in (m-r) to a binary phase map. Excitation NA: NA_od_=0.09, NA_id_=0.03.

We evaluated the intensity profile of the discontinuous light sheets created by scanning the same non-coaxial beam array synchronized with regional virtual confocal slits of different widths to understand whether the proposed method could work as expected to reject the off-focus fluorescence and how efficient the off-focus fluorescence is blocked at different confocal slit widths. We compared the intensity profile and thickness of the discontinuous light sheets obtained by scanning the non-coaxial beam array in Figure 5(a) synchronized with the regional virtual confocal slits in Figure 5(b) of various widths. The results (Figure 5(c)-5(l)) show that the regional virtual confocal slits reject the off-focus fluorescence background efficiently until the slit width reaches 120 μm, which is about 1.5 times of the beam array gap distance.

**Figure 5.**
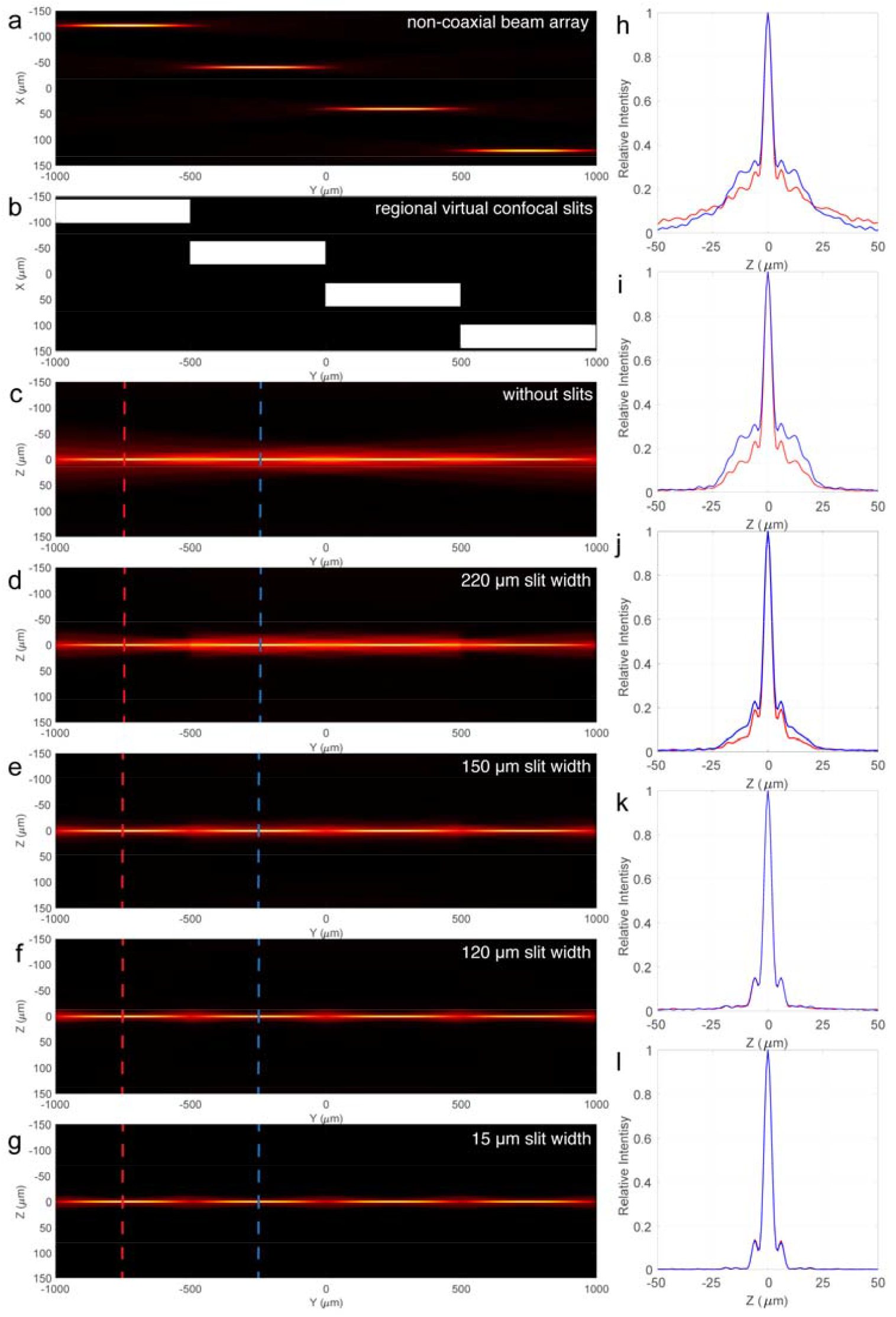
The ability of regional virtual confocal slits of different widths in rejecting the off-focus fluorescence generated by a non-coaxial beam array. (a) XY MIP of a four-beam non-coaxial beam array with 500 μm period and 80 μm gap distance. Excitation NA: NA_od_=0.06, NA_id_=0.02. (b) Regional virtual confocal slits with variable widths (X direction). (c-g) YZ MIPs of the discontinuous light sheets obtained by scanning the non-coaxial beam array in (a) symphonized with the regional virtual confocal slits of various widths in (b). (h-l) The intensity profile of the discontinuous light sheets in (c-g) at the indicated positions.

Next, we studied the influence of the beam array gap distance on the intensity profile of the discontinuous light sheets obtained by the synchronized scan of the non-coaxial beam array and regional virtual confocal slits to understand if narrower slits are needed to reject the off-focus fluorescence generated by non-coaxial beam arrays with short gap distances efficiently (Figure 6). We compared the intensity profile and thickness of the discontinuous light sheets obtained by scanning the non-coaxial beam array in Figure 6(a), which consists of the same excitation beams as that in Figure 5(a) but with a half of the gap distance, synchronized with the regional confocal slits in Figure 6(b) of various widths. Again, the results (Figure 6(c)-6(l)) show that the regional virtual confocal slits reject the off-focus fluorescence efficiently until the slit width reaches about 1.5 times of the beam array gap distance, which is about 60 μm. Therefore, we conclude that narrower slits are needed to block the off-focus fluorescence generated by non-coaxial beam arrays with smaller gap distances efficiently. Obviously, the synchronization of scanning non-coaxial beam arrays with larger gap distances and regional confocal slits with larger widths is easier and more robust although the imaging time is increased a little due to the longer scanning distance of the beam array. On the other hand, 3D imaging using regular continuous light sheets in TLSM can be considered as imaging using non-coaxial beam arrays with the gap distance larger than the height of the detection camera FOV.

**Figure 6.**
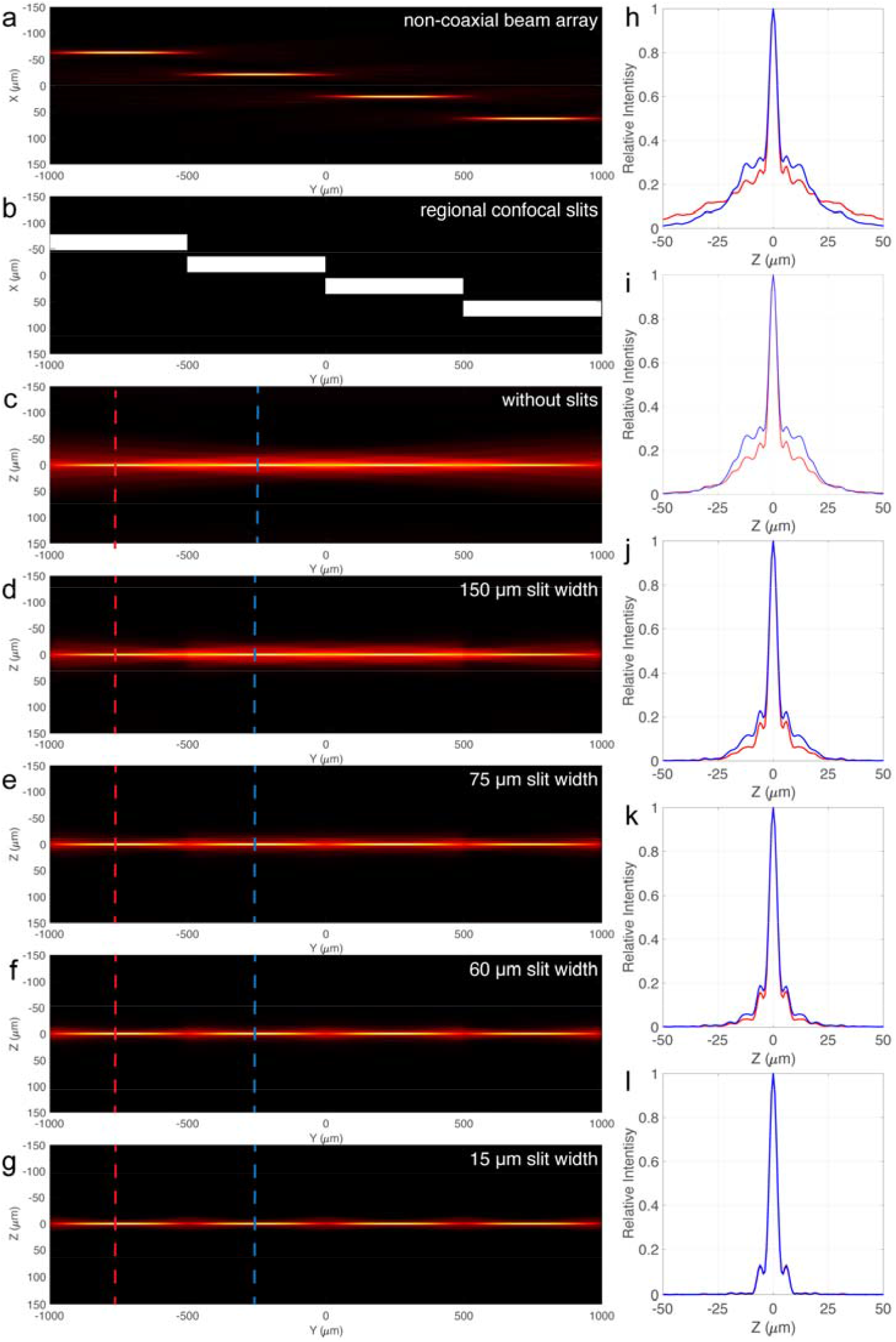
The maximal width of regional virtual confocal slits to reject the off-focus fluorescence generated by non-coaxial beam arrays efficiently decreases with the gap distance. (a) XY MIP of a four-beam non-coaxial beam array with 500 μm period and 40 μm gap distance. Excitation NA: NA_od_=0.06, NA_id_=0.02. (b) Regional virtual confocal slits with variable widths. (c-g) YZ MIPs of the discontinuous light sheets obtained by scanning the non-coaxial beam array in (a) symphonized with the regional virtual confocal slits of various widths in (b). (h-l) The intensity profile of the discontinuous light sheets in (c-g) at the indicated positions.

We further investigated whether narrower regional confocal slits are needed to block the off-focus fluorescence generated by non-coaxial beam arrays that consist of thinner excitation beams. We evaluated the intensity profile of the discontinuous light sheets obtained by synchronized scan of the non-coaxial beam array in Figure 7(a) with the regional confocal slits in Figure 7(b) of various widths, in which the non-coaxial beam array consists of thinner excitation beams than that of the non-coaxial beam array in Figure 5(a), but has the same beam number, gap distance and beam array period. The results show that the regional confocal slits also block the off-focus fluorescence background efficiently until the width of the slits reaches roughly 1.5 times of the beam array gap distance despite the slightly higher background, which suggests the off-focus fluorescence rejection efficiency by reginal confocal slits is unrelated to the intensity profile of individual excitation beams within the non-coaxial beam array.

**Figure 7.**
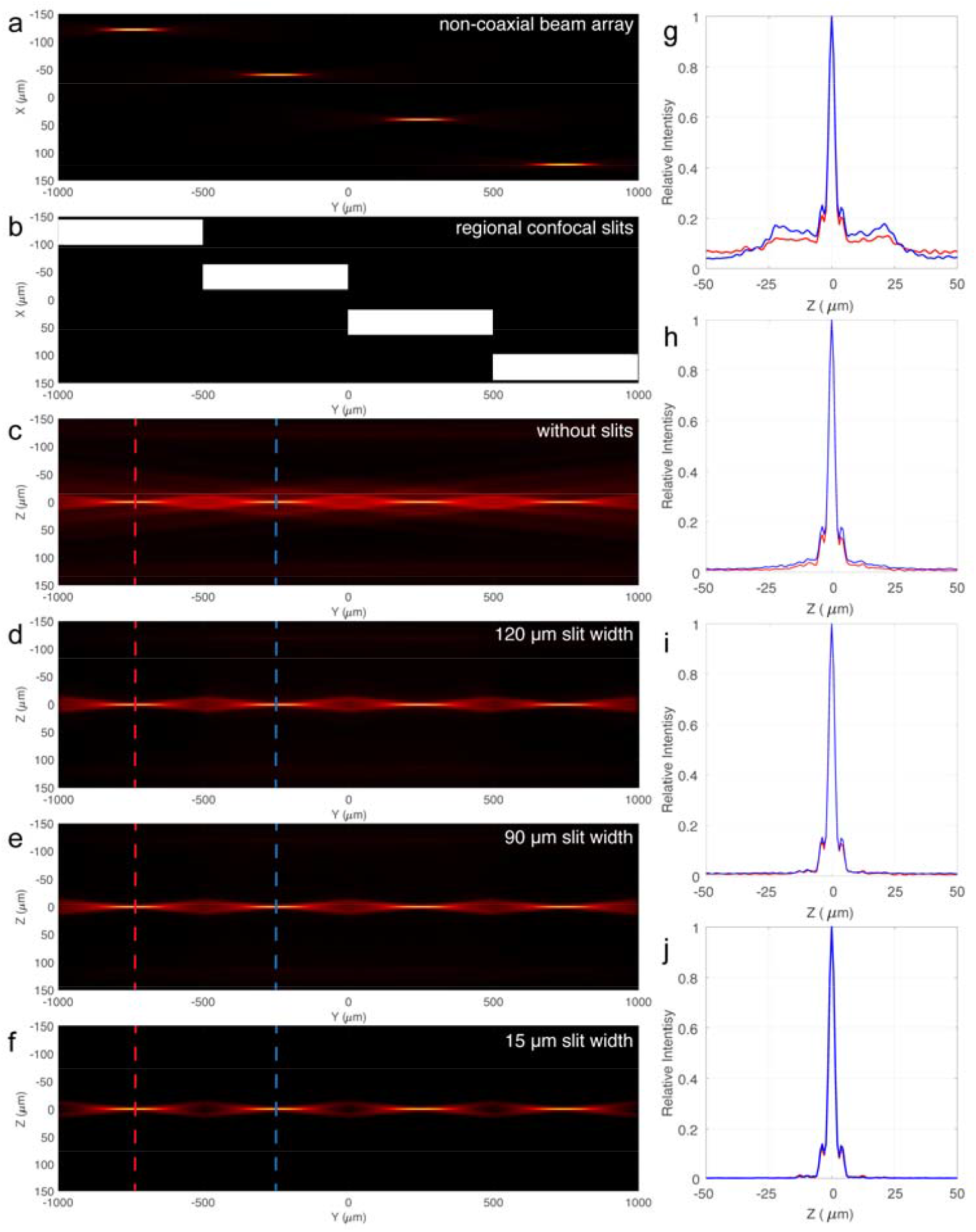
The efficiency of regional virtual confocal slits in reject the off-focus fluorescence generated by non-coaxial beam arrays is barely affected by the intensity profile of individual excitation beams. (a) XY MIP of a four-beam non-coaxial beam array with 500 μm period and 80 μm gap distance. Excitation NA: NA_od_=0.09, NA_id_=0.03. (b) Regional virtual confocal slits with variable widths. (c-f) YZ MIPs of the discontinuous light sheets obtained by scanning the non-coaxial beam array in (a) symphonized with the regional virtual confocal slits of various widths in (b). (h-l) The intensity profile of the discontinuous light sheets in (c-g) at the indicated positions.

## 4. Implementation

The realization of regional virtual confocal slits is critical to implement the proposed method in practice. In conventional sCMOS cameras, a global rolling shutter is used to control the exposure of the camera pixels. Different rows of camera pixels are exposed sequentially at different times as the rolling shutter sweeps through the camera sensor. As shown in Figure 8, the usage of a global rolling shutter produces a detection effect that is equivalent to the use a sweeping global virtual confocal slit. The sweeping speed of the global virtual confocal slit can be adjusted by changing the shift time of the rolling shutter. Both the shift time of the rolling shutter and the exposure time of each pixel row determine the number of pixel rows exposing at the same time, which define the width of the global virtual confocal slit. For instance, the sweeping speed of the virtual confocal slit can be increased by decreasing the rolling shutter shift time. The width of the global virtual confocal slit can be decreased by either increasing the rolling shutter shift time or decreasing the exposure time of each pixel row, so that less pixel rows are exposed at the same time which produces a narrower global virtual confocal slit.

**Figure 8.**
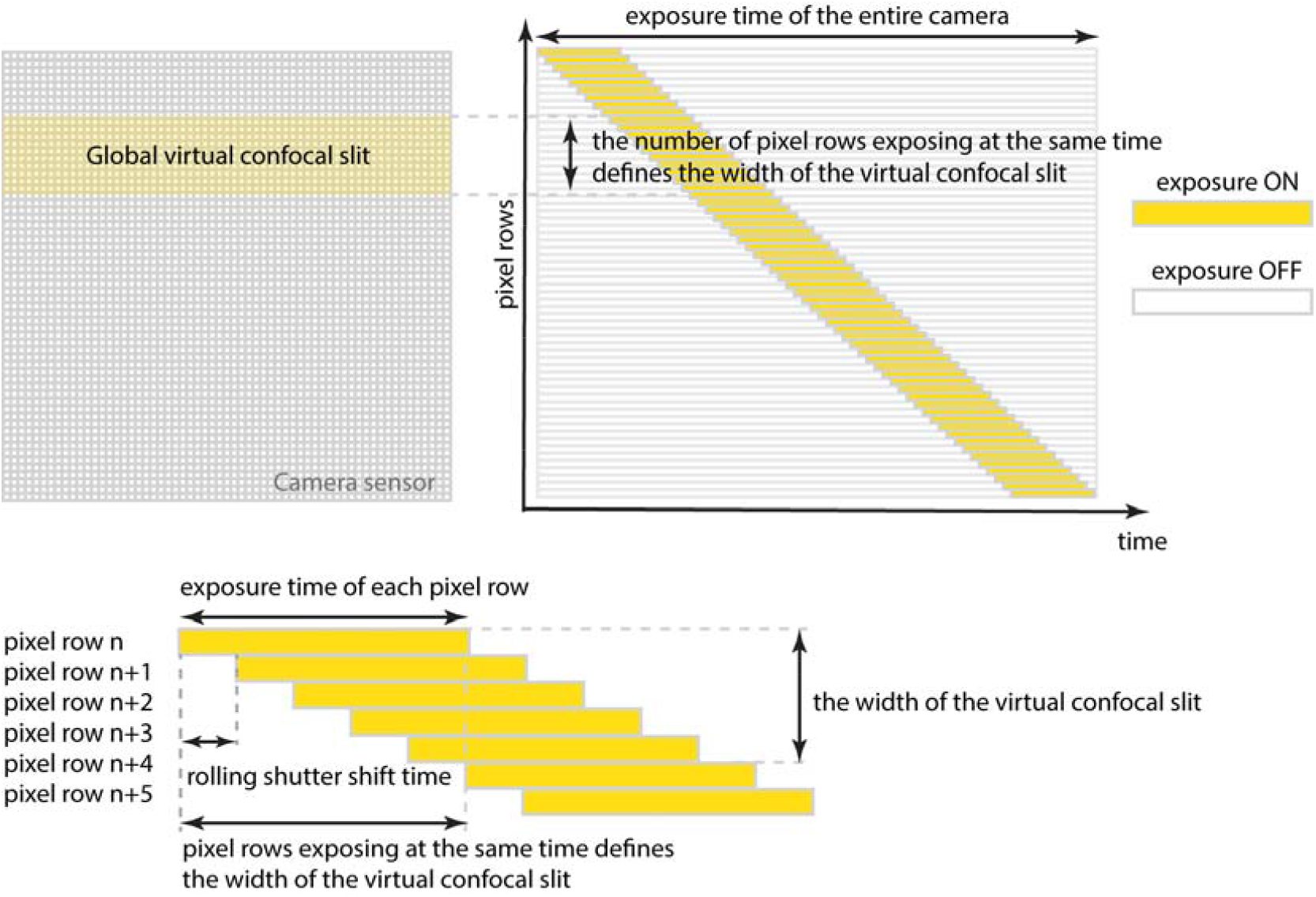
The working mechanism of the global virtual confocal slit that is controlled by the global rolling shutter of the detection camera.

Likewise, regional virtual confocal slits can be obtained easily by using a detection camera adopting multiple rolling shutters that control the exposure of different pixel rows in different camera regions separately. As shown in Figure 9, the detection camera is separated to four column regions, and the exposure of the four regions are controlled by four rolling shutters separately, which results in the realization of four regional virtual confocal slits. Besides the sweeping speed and width of each regional confocal slit are controlled by the corresponding rolling shutter individually, the gap distance between the adjacent regional virtual confocal slits is controlled by both the difference of the exposure initiation time and the shift time of different rolling shutters.

**Figure 9.**
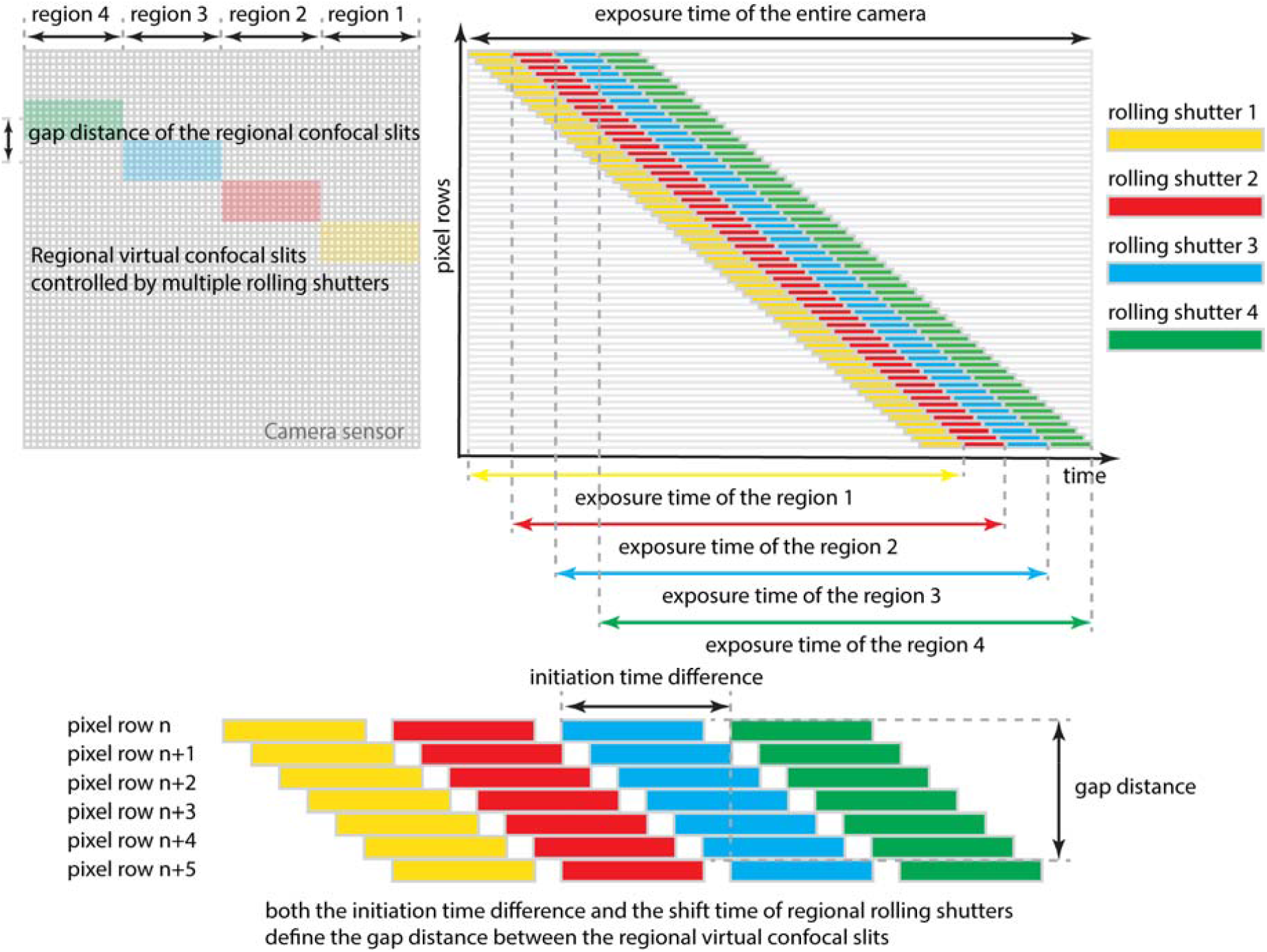
The realization of regional virtual confocal slits by adopting multiple rolling shutters that control different regional virtual confocal slits separately.

Obviously, the proposed method can be realized directly by using a scanning non-coaxial beam array synchronized with the regional virtual confocal slits on a sCMOS camera equipped with multiple rolling shutters (Figure 10(a)). The higher the number of rolling shutters, the more regional virtual confocal slits can be obtained, and the more excitation beams can be included in a non-coaxial beam array to improve the imaging efficiency of TLSM. Unfortunately, sCMOS cameras equipped with multiple rolling shutters are not yet available to our knowledge. However, there seems to be no technique barriers to prevent the development of such cameras, which would significantly advance the efficiency of TLSM for imaging cleared tissues as suggested by our results.

**Figure 10.**
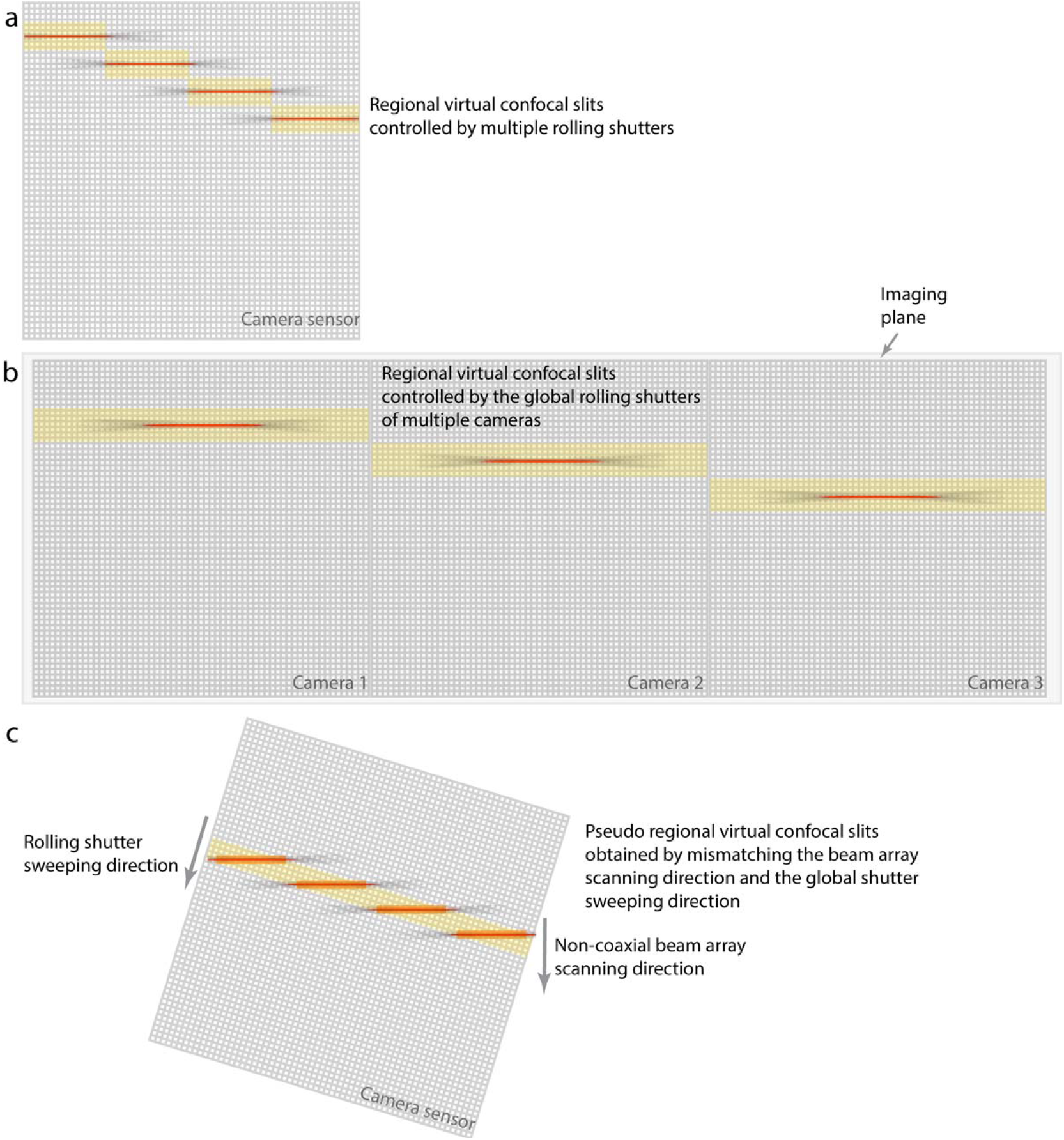
The implementation of regional confocal slits in practice. (a) A signal detection camera that is capable to control the camera exposure with multiple rolling shutters. (b) The global rolling shutters of multiple detection cameras. (c) Mismatch the non-coaxial beam array scanning direction and the rolling shutter sweeping direction.

Due to the lack of the camera described above, we propose two configurations to implement pseudo regional virtual confocal slits in practice. First, it is possible to use multiple sSMOS cameras for image detection (Figure 10(b)), so that the global rolling shutter of each camera controls one of the pseudo regional virtual confocal slits. Nevertheless, the optical configuration of the microscope is more complex to implement multiple detection cameras and there could be other physical restrictions limit the number of cameras can be used. Second, pseudo regional virtual confocal slits can also be obtained by mismatching the non-coaxial beam array scanning direction and the rolling shutter sweeping direction of a sCMOS camera (Figure 10(c)), so that the different regions within a single global virtual confocal slit can work as multiple regional virtual confocal slits due to the mismatching. However, this method also has many limitations. First, only a part of the camera can be used for imaging due to the scanning direction mismatch, which decreases the imaging efficiency. Second, the intensity profile of the non-coaxial beam array, including the beam number, gap distance and beam array period, is restrained by the mismatch angle between the beam array scanning direction and the rolling shutter sweeping direction. Third, the effective width of the regional virtual confocal slits created by this method is also limited by the mismatching angle and the non-coaxial beam array being used, which not only decreases the imaging efficiency and flexibility but also make the synchronization less robust.

## 5. Conclusions

In summary, we described a method of using scanning non-coaxial beam arrays synchronized with regional virtual confocal slits to improve the imaging efficiency of TLSM. We investigated the method to generate non-coaxial beam arrays using a SLM by applied phase maps calculated using a pupil segmentation method, characterized the method via numerical simulations, and we show the presented method could increase the imaging efficiency and feasibility of TLSM significantly. We also proposed several configurations to implement the method in practice. In addition, we describe a new detection camera operation mode that adopts multiple rolling shutters to control the exposure of camera pixel rows in different regions and enable the use of regional virtual confocal slits.

## Notes

### Competing Interest Statement

The authors have declared no competing interest.

